## Supplementary material for "Incorporating physics to overcome data scarcity in predictive modeling of protein function: a case study of BK channels": SI Figures and Tables

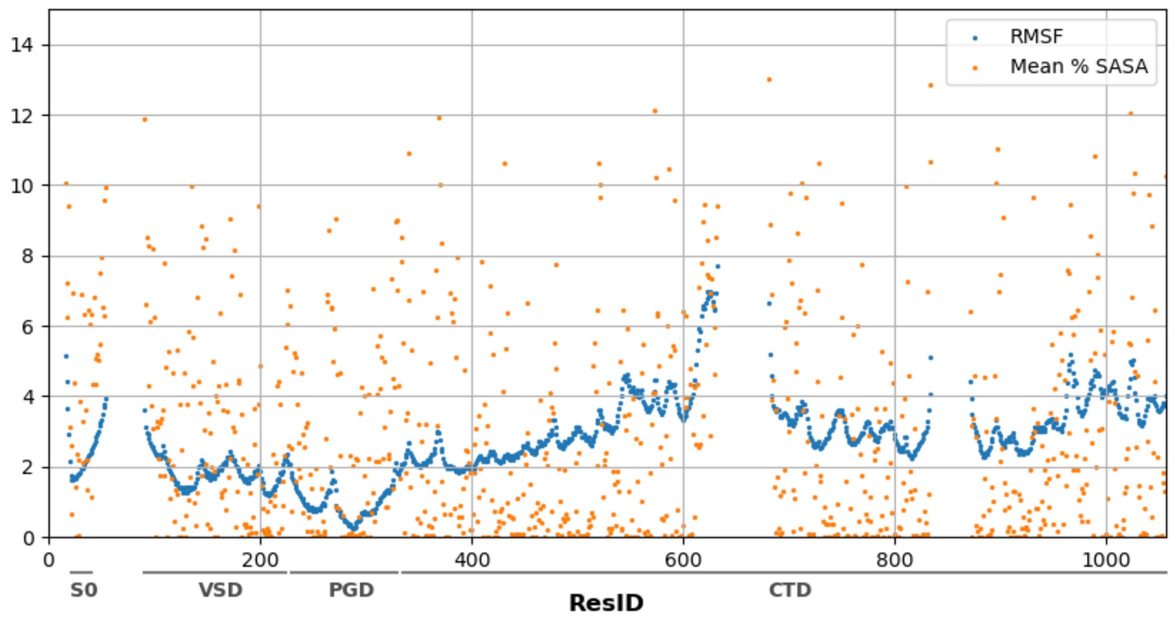


**Figure S1: Mean RMSF (blue) and percent SASA (orange) by Residue ID (ResID).** The RMSF has the units of Å, and the mean percent SASA is a unitless percentage. These features were calculated from 100 ns MD simulation of the deactivated BK channel.

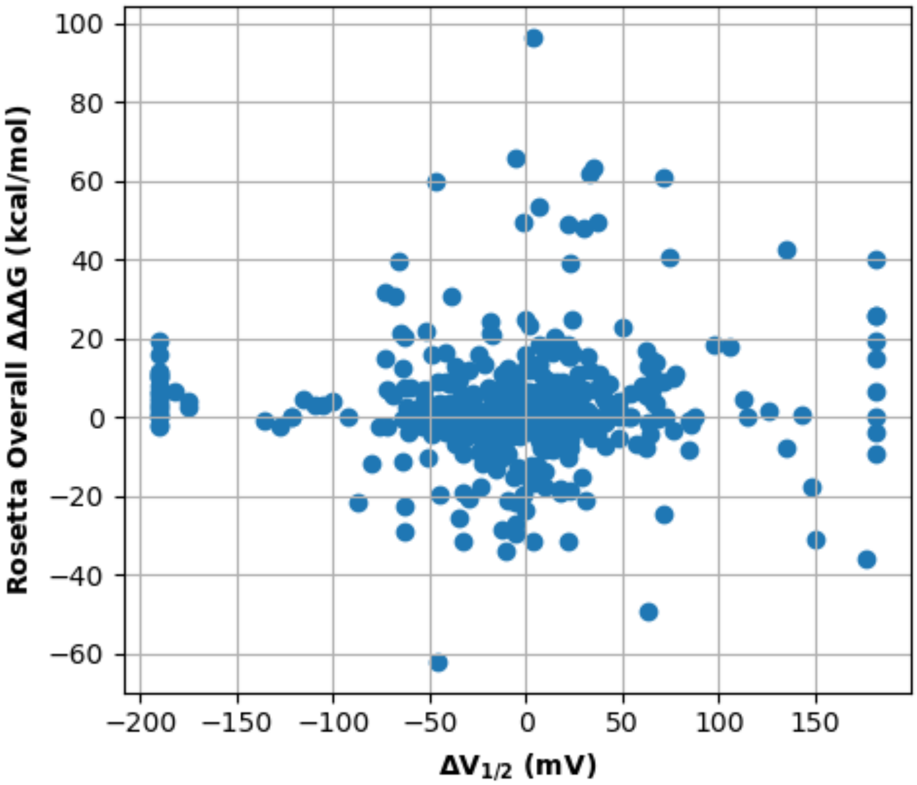


**Figure S2: Correlation of experimental $\Delta V_{1/2}$ with the total Rosetta ΔΔΔG.** The Pearson correlation coefficient is $R < 0.01$. See method section for description of how the Rosetta ΔΔΔG scores were calculated.

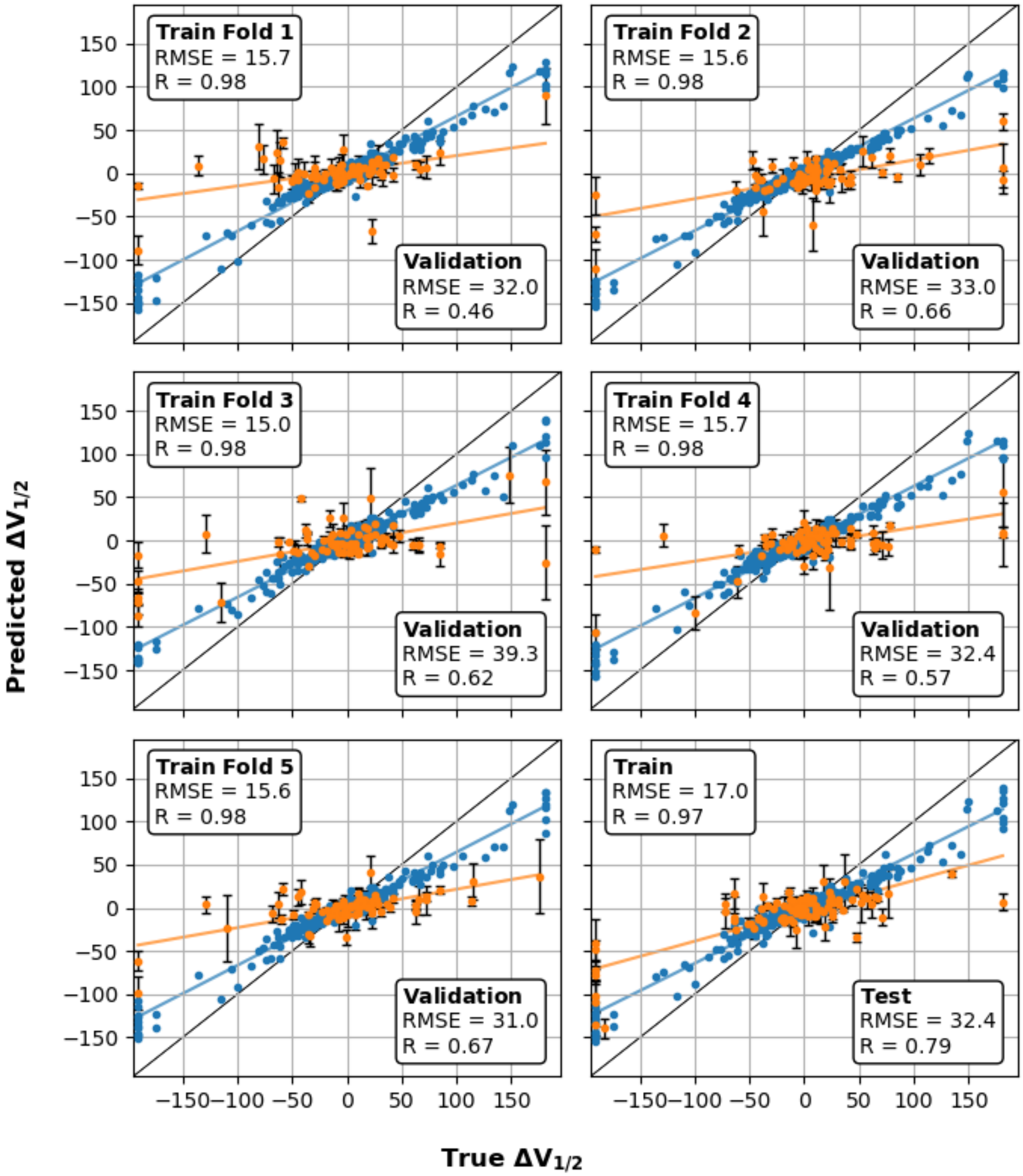


**Figure S3: 5-fold cross-validation on initial training split.** In all panels, blue denotes a data point used to train that iteration of the model, orange denotes validation or testing. The first five panels correspond to the 5-fold cross-validation within the training set (80% of the total data). The final panel (bottom right) was trained on the full training set and validated using the rest 20% of the full data set. This split corresponds to the 80/20 training/test split 1 from Table 1.

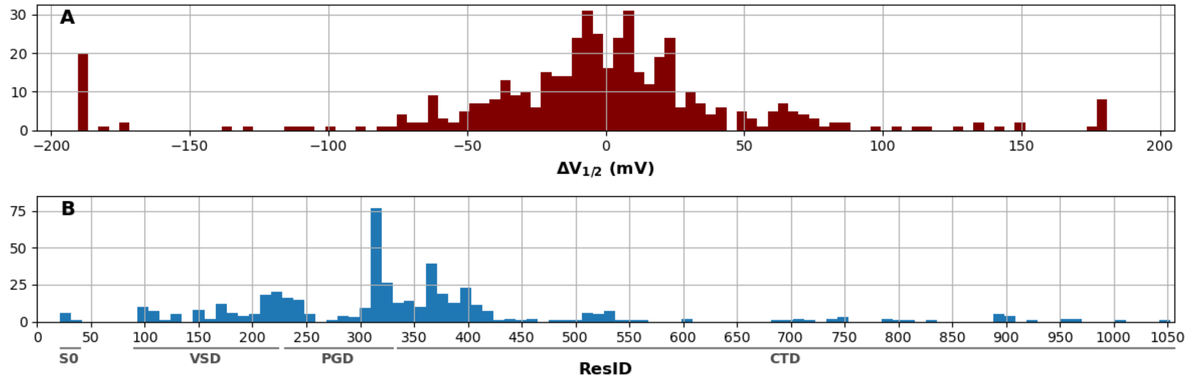


**Figure S4: Distributions of (A) experimental $\Delta V_{1/2}$ values and (B) number of existing mutations along the sequence.** Each histogram was generated using 100 bins. **(A)** depicts the $\Delta V_{1/2}$ distribution after squashing between ± 200 mV (see Methods). The key functional domains are highlighted beneath **(B)**.

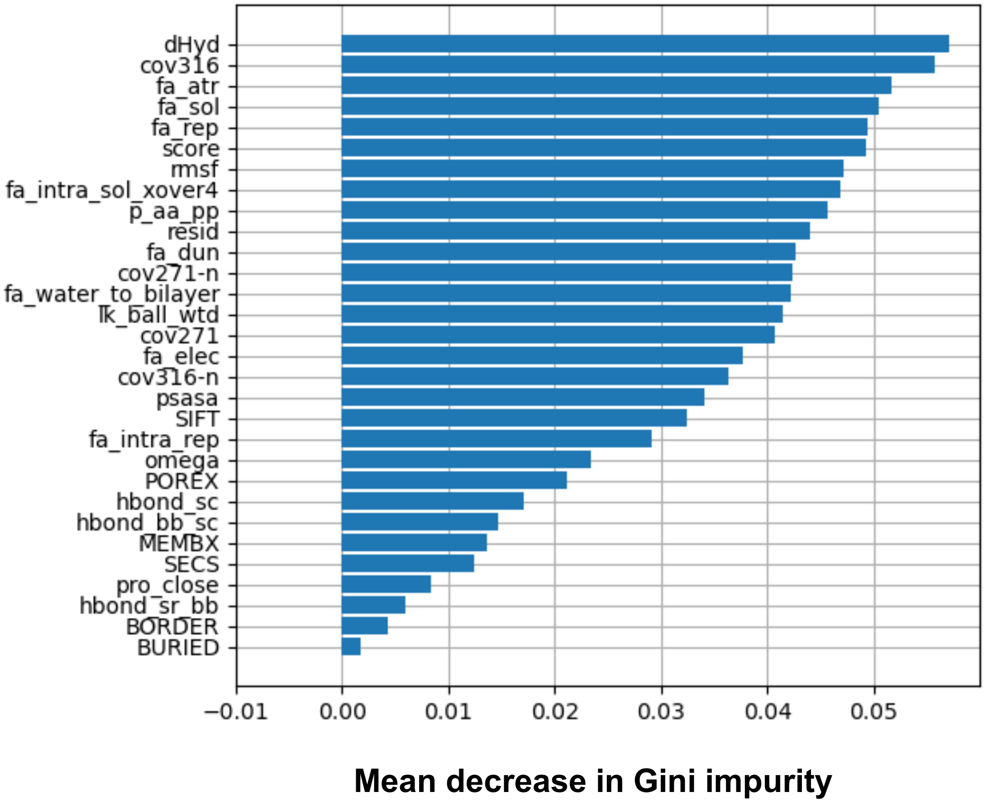


**Figure S5: Feature Importance.** Importance is reported as the mean decrease in Gini impurity score. A larger decrease in this impurity score means that using the feature in a branch of a tree often leads to greater ability to resolve gating voltage shifts. Names of features and their source are provided in Table S2.

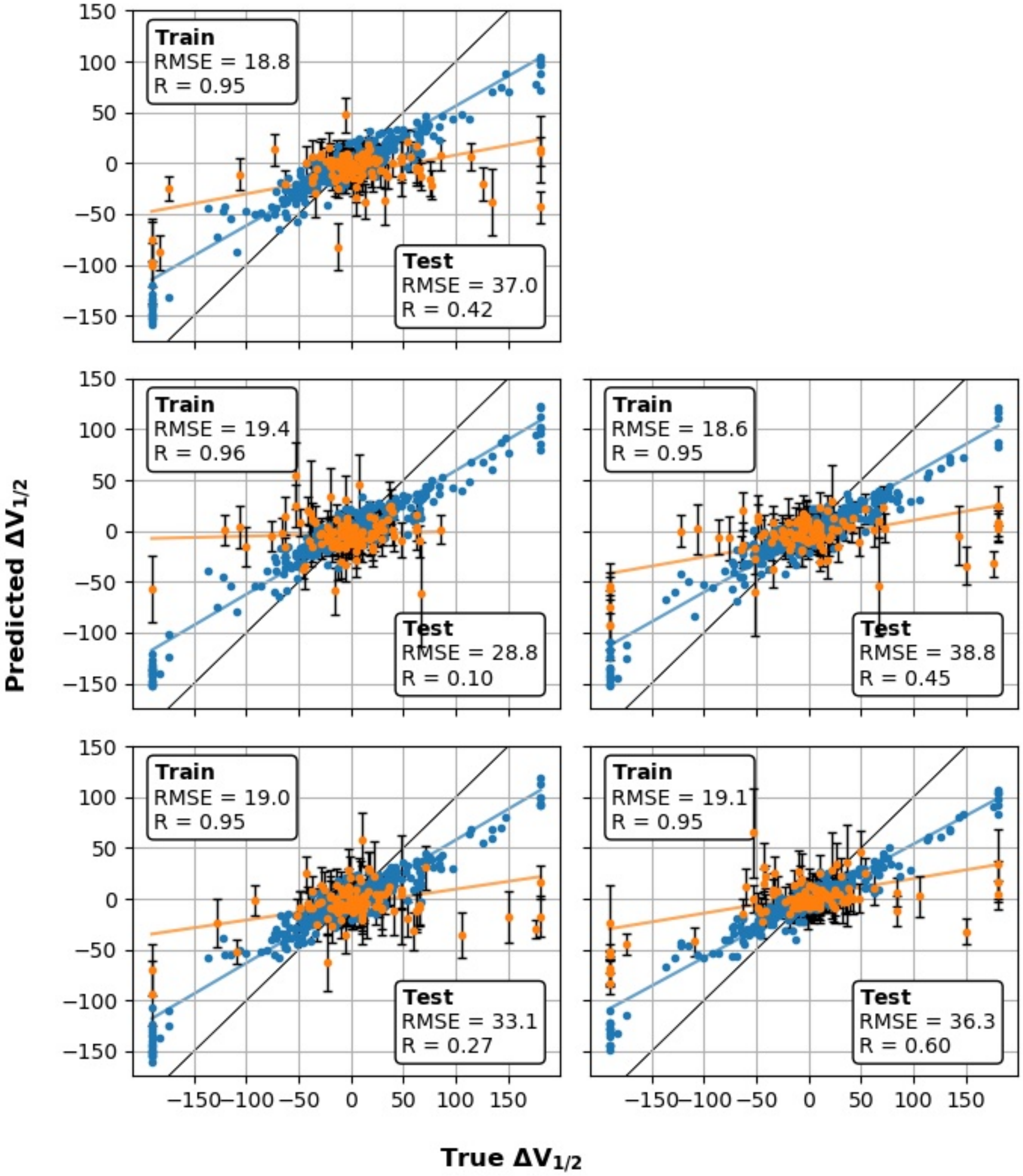


**Figure S6: Performance of control model trained without physics-based descriptors.** In all panels, blue dots denote data points used to train that iteration of the model, and orange dots denote the independent test data.

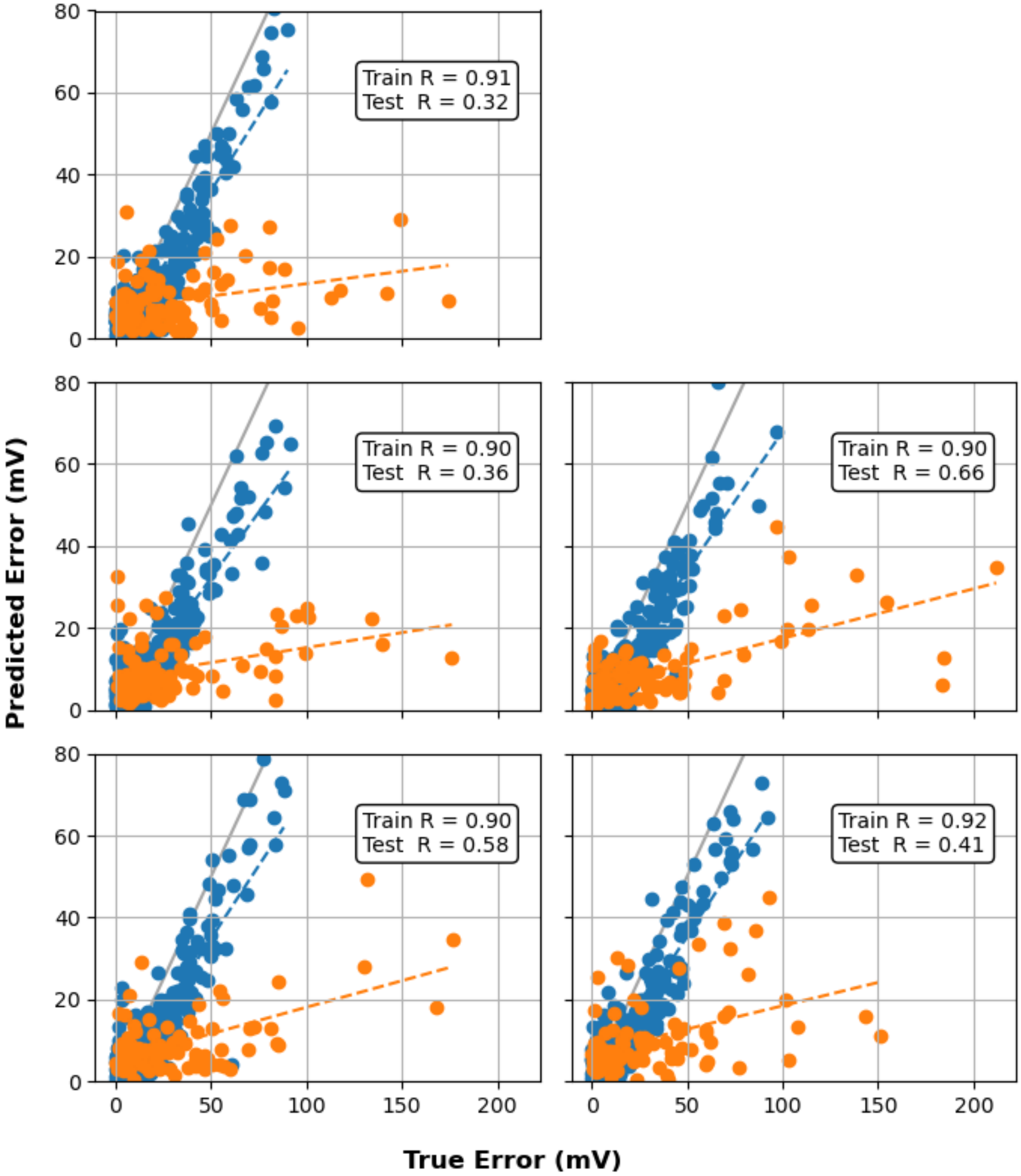


**Figure S7: Correlation of the true and predicted errors for the train and test sets of the five independent data splits.** Blue and orange dot represent training and test data, respectively. The dashed lines denote lines of best fit with the same color scheme. The gray lines denote y=x.

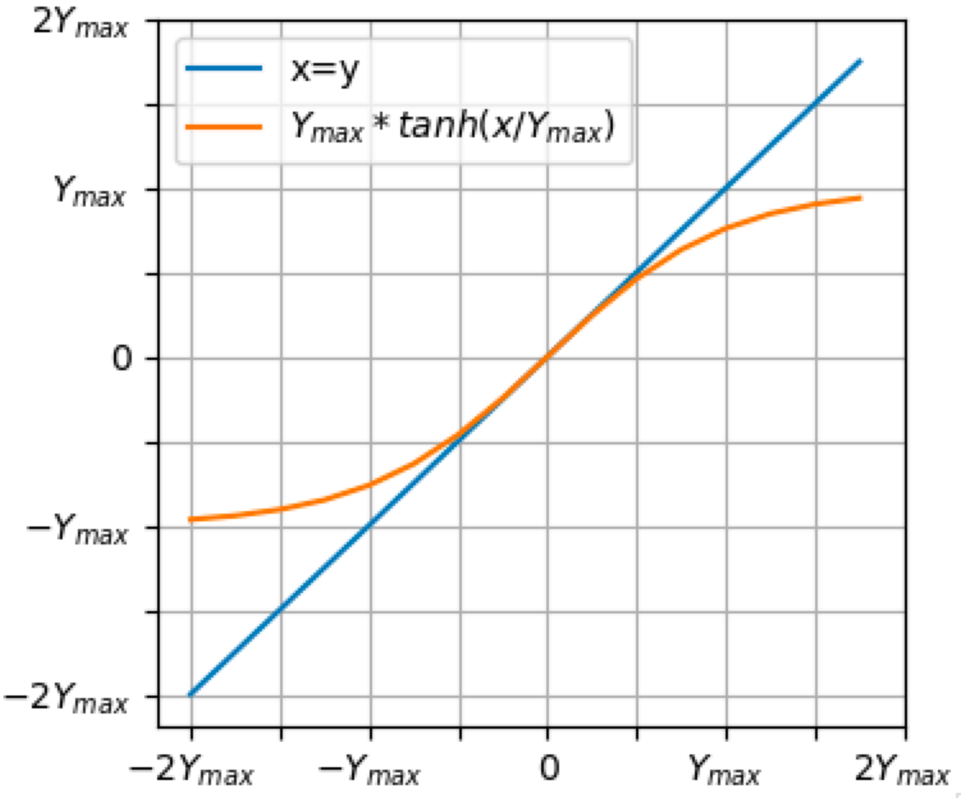


**Figure S8: Illustration of the squashing function for pre-processing of various raw quantities.** The blue line plots the original, un-squashed function as a reference.

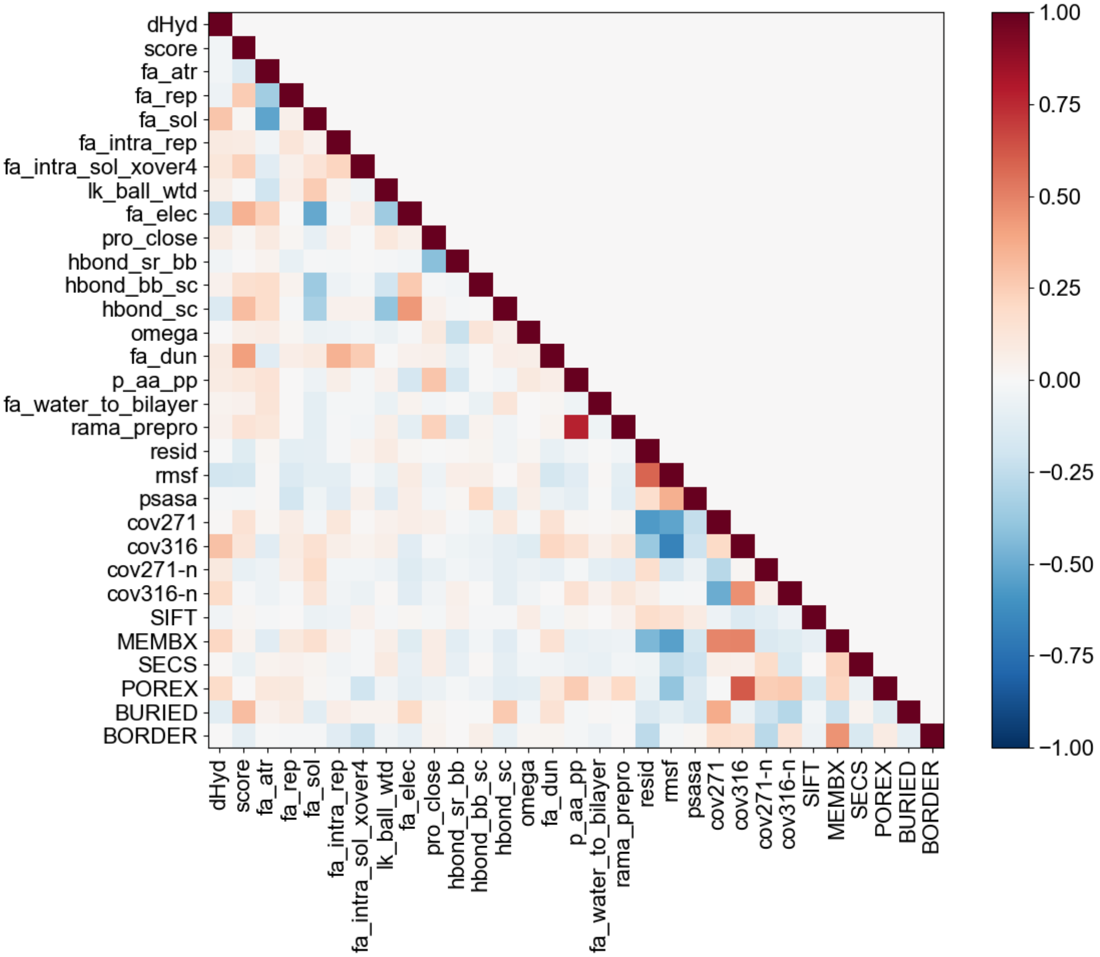


**Figure S9: Feature correlations.** Correlation of the features used in the model training. Correlation is reported as the Pearson correlation coefficient. Names and descriptions of features and their sources are provided in Table S1.

**Table S1: List of features and their descriptions.** All Rosetta terms are ΔΔΔG values between the closed and open states as described in Methods.

| Feature[#] | Description[#] | Method |
|---|---|---|
| score | Total $\Delta G_{fold}$ | Rosetta ΔΔΔG |
| fa_atr | Lennard-Jones attractive between atoms in different residues. | Rosetta ΔΔΔG |
| fa_rep | Lennard-Jones repulsive between atoms in different residues. | Rosetta ΔΔΔG |
| fa_sol | Lazaridis-Karplus solvation energy. | Rosetta ΔΔΔG |
| fa_intra_rep | Lennard-Jones repulsive between atoms in the same residue. | Rosetta ΔΔΔG |
| fa_elec | Coulombic electrostatic potential with a distance-dependent dielectric. | Rosetta ΔΔΔG |
| pro_close | Proline ring closure energy and energy of Ψ angle of preceding residue. | Rosetta ΔΔΔG |
| hbond_sr_bb | Backbone-backbone hydrogen bonds close in primary sequence. | Rosetta ΔΔΔG |
| hbond_lr_bb | Backbone-backbone hydrogen bonds distant in primary sequence. | Rosetta ΔΔΔG |
| hbond_bb_sc | Sidechain-backbone hydrogen bond energy. | Rosetta ΔΔΔG |
| hbond_sc | Sidechain-sidechain hydrogen bond energy. | Rosetta ΔΔΔG |
| dslf_fa13 | Disulfide geometry potential. | Rosetta ΔΔΔG |
| rama | Ramachandran preferences. | Rosetta ΔΔΔG |
| omega | A harmonic restraint on planarity of the ω backbone dihedral with standard deviation of ~6°. | Rosetta ΔΔΔG |
| fa_dun | Internal energy of sidechain rotamers as derived from Dunbrack's rotamer statistics (2010 Rotamer Library used in Talaris2013). | Rosetta ΔΔΔG |
| p_aa_pp | Probability of amino acid at Φ/Ψ. | Rosetta ΔΔΔG |
| yhh_planarity | A special torsional potential to keep the tyrosine hydroxyl in the plane of the aromatic ring. | Rosetta ΔΔΔG |
| ref | Reference energy for each amino acid that balances the internal energy of amino acid terms. | Rosetta ΔΔΔG |
| fa_intra_sol_xover4 | Intra-residue LK solvation, counted for the atom-pairs beyond torsion-relationship. | Rosetta ΔΔΔG |
| fa_intra_elec | Intra-residue Coulombic interaction, counted for the atom-pairs beyond torsion-relationship. | Rosetta ΔΔΔG |
| rama_prepro | Backbone torsion preference term that accounts for whether preceding amino acid is Proline or not. | Rosetta ΔΔΔG |
| lk_ball | Anisotropic contribution to the solvation. | Rosetta ΔΔΔG |

| | | |
|---|---|---|
| cov316 | Row of intra-monomer Ca-Ca covariance matrix corresponding to residue A316, describes coupled motion with the pore | MD |
| cov278 | Row of intra-monomer Ca-Ca covariance matrix corresponding to residue X278, 278 also has several mutations with large shifts and is not covariant with the pore | MD |
| cov316-n | Row of directly-neighboring inter-monomer Ca-Ca covariance matrix corresponding to residue A316, describes coupled motion with the pore | MD |
| cov278-n | Row of directly-neighboring inter-monomer Ca-Ca covariance matrix corresponding to residue X278, 278 also has several mutations with large shifts and is not covariant with the pore | MD |
| RMSF | Root-mean square fluctuations, describes chain flexibility | MD |
| pSASA | Percent Solvent-Accessible Surface Area, describes solvation effects including in the deep pore. | MD |
| dHyd | Change in hydrophobicity from wildtype to mutant. Hydrophobicity is transfer free energy from cyclohexane to water. | Biochemical data (1) |
| ResID | The residue ID or number, following the numbering scheme of the PDB entries. | -- |
| WATX | Residue is within 5 Å of water in equilibrated snapshot | MD |
| PORX | Residue is pore-lining | Structure |
| MEMX | Residue is within 5 Å of lipid in equilibrated snapshot | Structure |
| BORDER | Both WATX and MEMX, residue is at membrane interface | MD/Structure |
| SECS | Secondary Structure (helix, coil, sheet) | Structure |
| SIFT | Score from the SIFT algorithm: Sorting Intolerant from Tolerant, predict effect of single nucleotide polymorphism based partially on sequence conservation score | SIFT tool (2) |

[#] Based on: https://www.rosettacommons.org/docs/latest/rosetta_basics/scoring/score-types and https://www.ncbi.nlm.nih.gov/pmc/articles/PMC5717763/table/T1/

**Table S2: Summary of machine learning models, hyperparameters trained in Grid Search, and training and validation correlation (R) and RMSE from 5 independent splits of data.**

| Model | Hyperparameters | R (Train, Test) | RMSE (mV) (Train, Test) |
|---|---|---|---|
| Ridge | **alpha**: 1e-6, 1e-4, 1e-2, 1, 10, **1e2**, 1e4, 1e6 | (0.40, 0.33), (0.39, 0.29), (0.41, 0.21), (0.39, 0.22), (0.41, 0.24) | (37, 44), (39, 37), (37, 44), (37, 44), (38, 42) |
| SVR | **kernel**: 'linear', 'poly', 'sigmoid', **'rbf'**; **degree**: 2, **3**, 6; **C**: 1e-2, 1e-1, 1, 1e1, **1e2**; **coef0**: 0, **1**; **epsilon**: 0, 1e-4, 1e-2, **1**, 1e2 | (0.81, 0.69), (0.83, 0.52), (0.83, 0.57), (0.80, 0.60), (0.82, 0.57) | (19, 35), (18, 32), (18, 36), (18, 34), (18, 35) |
| RF | **n_estimators**: 50, 100, 250, **500**, 1000; **min_samples_split**: **2**, 5, 10, 20; **max_leaf_nodes**: 2, 5, 10, 20, 50, **100**, 200; **max_depth**: 2, 5, 10, **20**, 50, 100; **max_features**: 0.1, 0.25, 0.5, 0.75, 0.9, 1.0; **ccp_alpha**: 1e-3, **1e-2**, 1e-1, 1, 10, 100; **max_samples**: 0.1, 0.25, 0.5, 0.75, **1.0**; **min_samples_leaf**: **1**, 2, 5, 10, 20; **min_weight_fraction_leaf**: **0**, 0.01, 0.1, 0.25, 0.5; | (0.97, 0.79), (0.97, 0.54), (0.97, 0.69), (0.97, 0.80), (0.98, 0.70) | (17, 32), (17, 30), (16, 35), (17, 31), (16, 31) |
| KNN | **n_neighbors**: **3**, 7, 11, 21, 51, 101; **weights**: **'uniform'**, 'distance'; **leaf_size**: **5**, 10, 50, 100; **p**: **1**, 2, 3; **algorithm**: 'ball_tree', **'kd_tree'**, 'brute' | (0.81, 0.73), (0.84, 0.62), (0.83, 0.77), (0.82, 0.63), (0.82, 0.52) | (23, 30), (21, 29), (22, 29), (23, 33), (22, 34) |
| GP | **n_estimators**: 10, **50**, 100, 500, 1000; **max_depth**: 2, 4, 16, **32**, 64; **min_samples_split**: 2, 4, 16, 32, 64; **max_features**: 0.25, 0.5, **0.75**, 1.0; **ccp_alpha**: 0, **5e-3**, 10e-3, 20e-3 | (0.92, 0.57), (0.87, 0.30), (0.87, 0.54), (0.92, 0.62), (0.87, 0.64) | (34, 41), (36, 34), (34, 42), (35, 39), (35, 38) |
| MLP | **solver**: 'sgd'; **learning_rate**: invscaling; **power_**t: 1e-6, 1e-4, **1e-2**, 1; **activation**: 'tanh'; **hidden_layer_sizes**: (10), (50), **(100)**, (500), (100,10,100); **alpha**: 1e-6, 1e-4, 1e-2, **1** | (1.0, 0.77), (1.0, 0.40), (1.0, 0.76), (1.0, 0.74), (1.0, 0.66) | (2, 30), (2, 37), (2, 31), (2, 31), (2, 35) |

**Table S3: Neurological BK channel mutants, channel activity phenotypes, functional mechanisms (if known), and predicted $\Delta V_{1/2}$ values.** Abbreviations: VUS: Variant of Uncertain Significance, NE: No effect, LOF: Loss of Function, and GOF: Gain of Function. The Coordination of Rare Diseases at Sanford (CoRDS) is a standardized patient registry in a de-identified format. NP stands for No Prediction; the model doesn't predict any shift for mutations to residues absent in either the $Ca^{2+}$-bound or unbound PDB structure.

| PDB Mutation Name | Mutation Name (hslo1 gene) | BK channel activity | Functional Mechanism | Predicted $\Delta V_{1/2}$ (mV) | Reference |
|---|---|---|---|---|---|
| | G20D | VUS | | NP | (3) |
| | I29F | VUS | | NP | (3) |
| L206F | L271F | VUS | | 19 ± 14 | (3) |
| S286Y | S351Y | LOF | No current at 160 mV and 10 μM $Ca^{2+}$. | 14 ± 3 | (4) |
| G289S | G354S | LOF | G-V shift to depolarized potentials, restored by NS1619 | 5 ± 13 | (5) |
| G291R | G356R | LOF | No current at 160 mV and 10 μM $Ca^{2+}$. | -9 ± 12 | (4) |
| G310R | G375R | LOF | No Current | -2 ±7 | (4) |
| C348Y | C413Y | LOF | G-V shift to depolarized potentials, decreased expression | 5 ± 9 | (4) |
| D369G | D434G | GOF | G-V shift to hyperpolarized potentials, increased open probability, faster activation, slower deactivation, increased $Ca^{2+}$ sensitivity | -13 ± 4 | (5) |
| H379Q | H444Q | LOF | | 3 ± 4 | (6), CoRDS |
| K392E | K457E | LOF | G-V shift to depolarized potentials | 32 ± 12 | (7) |
| I447V | I512V | VUS | | -2 ± 2 | (8), CoRDS |
| K453N | K518N | NE | | -9 ± 1 | (9) |
| A467V | A532V | VUS | | -7 ± 4 | CoRDS |

| | | | | | |
|---|---|---|---|---|---|
| N471H | N536H | GOF | | -8 ± 5 | (10) |
| E500A | E565A | NE | | NP | (9) |
| G502S | G567S | VUS | | -3 ± 4 | CoRDS |
| E591A | E656A | VUS | | NP | (9) |
| I598V | I663V | LOF | | 3 ± 3 | (4) |
| E671K | E736K | VUS | | NP | (3) |
| D735Y | D800Y | VUS | | NP | CoRDS |
| P740L | P805L | LOF | G-V shift to depolarized potentials, decreased expression | -17 ± 5 | (4) |
| E819K | E884K | NE | | 4 ± 13 | (11) |
| D900V | D965V | LOF | | -7 ± 6 | (3, 6) |
| D919N | D984N | LOF | | NP | (4) |
| N988S | N1053S | GOF | G-V shift to hyperpolarized potentials, increased open probability, faster activation, slower deactivation, $Ca^{2+}$-independent mechanism | NP | (3), CoRDS |
| R1018K | R1083K | VUS | | -13 ± 5 | (12) |
| R1032H | R1097H | LOF | | NP | (6) |
| T1046R | T1111R | VUS | | -6 ± 7 | (3) |
| R1063W | R1128W | NE | | NP | (13) |
| T1089I | T1154I | VUS | | NP | (3) |
| N1094S | N1159S | NE | | NP | (9) |
